# Preserved functional connectivity in the default mode and salience networks is associated with youthful memory in superaging

**DOI:** 10.1101/254193

**Authors:** Jiahe Zhang, Joseph Andreano, Bradford C. Dickerson, Alexandra Touroutoglou, Lisa Feldman Barrett

## Abstract

“Superagers” are older adults who, despite their advanced age, maintain youthful memory. Previous morphometry studies revealed multiple default mode network (DMN) and salience network (SN) regions whose cortical thickness is preserved in superagers and correlates with memory performance. In this study, we examined the intrinsic functional connectivity within DMN and SN in 41 young (24.5 ± 3.6 years old) and 40 elderly adults (66.9 ± 5.5 years old). As in prior studies, superaging was defined as youthful performance on a memory recall task, the California Verbal Learning Test (CVLT). Participants underwent a resting state fMRI scan and performed a separate visual-verbal recognition memory task. As predicted, within both DMN and SN, superagers had stronger connectivity compared to typical older adults and similar connectivity compared to young adults. Superagers also performed similarly to young adults and better than typical older adults on the recognition task, demonstrating youthful episodic memory that generalized across memory tasks. Stronger connectivity within each network independently predicted better performance on both the CVLT and recognition task in older adults. Variation in intrinsic connectivity explained unique variance in memory performance, above and beyond preserved neuroanatomy. A post-hoc analysis revealed that DMN and SN nodes were more strongly inversely correlated in superagers than in typical older adults but were similarly correlated in superagers and young adults. Stronger between-network inverse correlations also predicted better memory performance in the entire sample of older adults. These results extend our understanding of the neural basis of superaging as a model of successful aging.

**SIGNIFICANCE STATEMENT:** Memory capacity generally declines with age, but a unique group of older adults – ‘superagers’ – have memory capacities rivaling those of younger adults, as well as preserved neuroanatomy in an ensemble of regions contained in two core intrinsic brain networks – the default mode and salience networks. In this study, we assessed the strength of intrinsic connectivity within these networks in superagers and typical older adults compared to young adults. We also expanded the behavioral assessment of memory. As predicted, superagers have intrinsic connectivity within the default mode and salience networks that is stronger than typical older adults and similar to that of young adults. Within older adults, preserved intrinsic connectivity within each network was uniquely associated with better memory performance.

---

Episodic memory decline is thought to accompany normal human aging (1, 2). Cross-sectional and longitudinal studies show that healthy older adults over the age of 70 perform less well on memory tests when compared to their younger counterparts and younger selves (3, 4). There is substantial individual variation in memory ability among the elderly, however, with some showing remarkable resilience to typical age-related cognitive decline (5-11). Recent research has identified a specific subgroup of elderly adults, called ‘superagers,’ whose recall memory on tests such as the Rey Auditory Verbal Learning Test or California Verbal Learning Test (CVLT) rivals that of middle-aged adults (12-15) and even young adults (16).

In addition to superior memory performance, superagers are further characterized by preserved neuroanatomy in an ensemble of brain regions that can be topographically found in two large-scale brain networks (16) – the default mode network (DMN; (17, 18)) and the salience network (SN; (19, 20)). The relevant regions and their network affiliations are presented in Fig. 1. These regions typically show age-related atrophy in healthy older adults (21-24).

**Figure 1.**
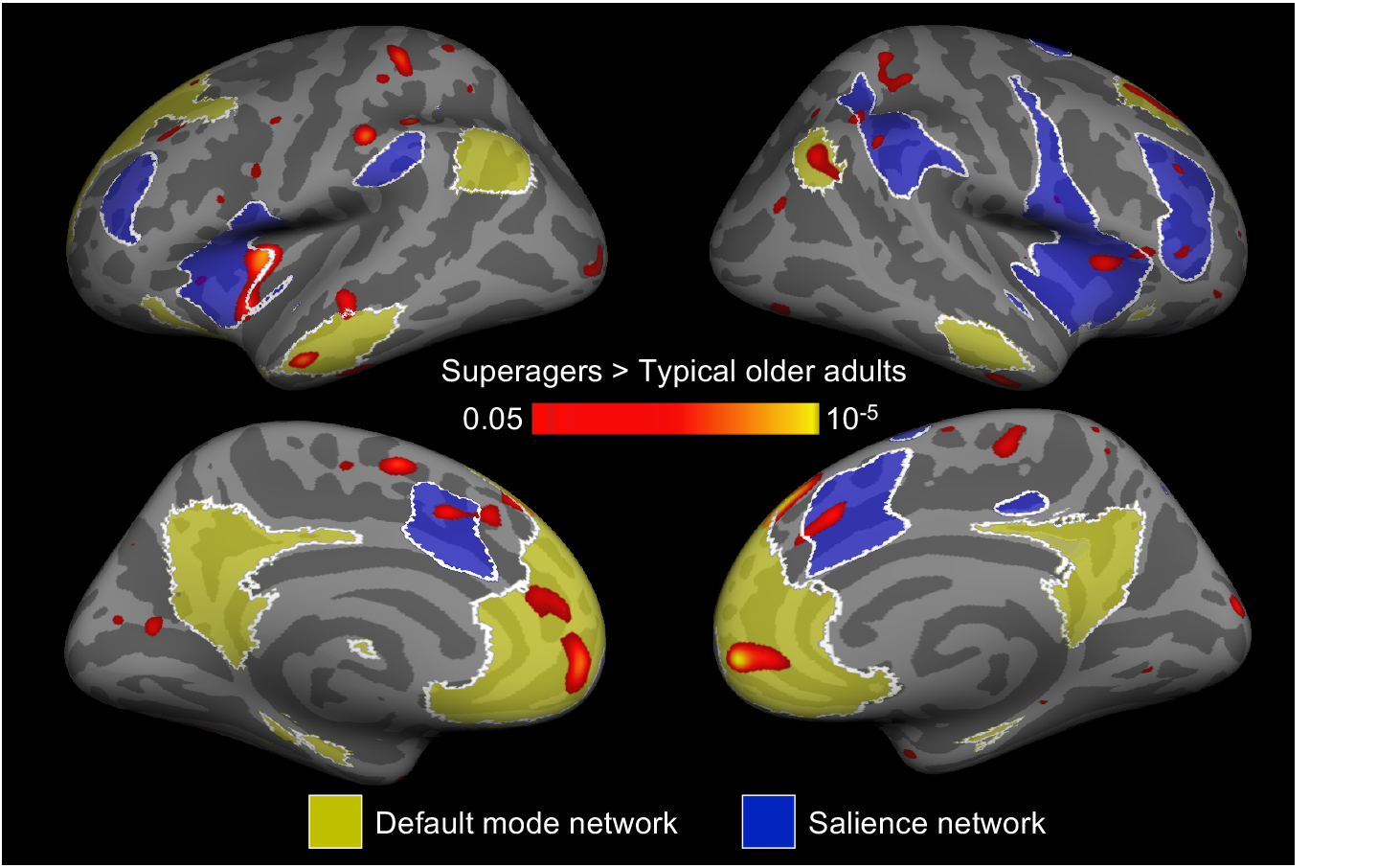
Regions of the DMN (yellow) and SN (blue) where superagers have thicker cortex than typical older adults. Map was thresholded at *p* < 0.05. Figure was adapted from (16).

Both the default mode and salience networks are implicated in many different psychological and physiological functions (25-28), including memory function (29, 30). Within the DMN, the hippocampal formation (HF) (31), medial temporal cortices, posterior cingulate cortex (PCC) and adjacent midline parietal areas (32) have all been implicated in the successful memory encoding and retrieval. Within the SN, main nodes are the anterior insula, anterior to mid cingulate cortex, as well as other frontoparietal regions including the middle frontal gyrus and inferior parietal lobule. The SN is broadly construed to be associated with attention (33) and executive function (34, 35), and several of these nodes are also shared with the frontoparietal control network (36, 37). The SN has been observed to support episodic memory encoding and retrieval (29, 38-40), possibly by directing attention to relevant material, engaging working memory, organizing available information and adjusting motivation (41, 42). In older adults, age-related atrophy in a number of DMN and SN regions is associated with memory decline (43-45), whereas their preservation predicts youthful verbal memory performance (16).

In addition to structural atrophy, typical aging is associated with progressively weaker intrinsic functional connectivity in the DMN (46-49) and SN (49-55). Reduced intrinsic connectivity between regions within the DMN has been linked to worse memory performance both in young (56) and in older adults (46, 47, 57). Reduced connectivity between regions within the SN has been associated with worse performance in attention and executive function tasks both in young (20) and in older adults (58), as well as with worse memory performance in young adults (Andreano et al., 2017). There is currently no evidence of a direct link between age-related decline in SN connectivity and memory in older adults, but one task-based functional MRI study has demonstrated that weaker changes in anterior cingulate cortex activity has been associated with poorer episodic memory in older adults (59).

In the current study, we built on these existing findings to examine the network coherence of the DMN and SN in superaging. We examined intrinsic functional connectivity of the DMN and SN in the same sample of healthy older adults (n = 40, 20 males, 66.9 ± 5.5 years old) and young adults (n = 41, 20 males, 24.5 ± 3.6 years old) as discussed in (16); these participants were part of a larger, longitudinal study, allowing us to report additional data and analyses that were not yet available at the time the initial report was submitted for publication. Participants completed an expanded set of episodic memory tests: as in previous studies, they completed the CVLT to assess verbal recall memory and to define superagers. Participants also viewed visual images (faces and scenes) paired with words to assess item and associative recognition memory. Prior to the recognition task, participants underwent a structural MRI scan and a resting state fMRI scan.

We hypothesized that superagers (i.e., individuals with CVLT scores comparable to young adults) would have preserved intrinsic connectivity within the DMN and SN relative to typical older adults but similar to that observed in young adults. We additionally hypothesized that superagers would show youthful memory performance on in both item and associative recognition, demonstrating that their preserved memory function generalized beyond their performance on the CVLT. Finally, we hypothesized that within older adults, individuals with stronger intrinsic DMN and SN connectivity would perform better on all memory tasks -- the verbal recall task (the CVLT) and the visual-verbal recognition task (for both item and associative memory). Given the different functional roles of the DMN and SN in memory, we hypothesized that DMN and SN connectivity would each independently predict memory performance relative to one another, as well as above and beyond the variance explained by the anatomical preservation of regions in those networks.

## RESULTS

### Default Mode and Salience Network Intrinsic Connectivity is Preserved in Superagers

The average intrinsic functional connectivity map for superagers and typical older adults can be found in Fig. S1 (for cortical regions) and Fig. 5A (for the hippocampus). The map for the DMN was anchored in a PCC seed as used in (36); the map for the SN was anchored in a seed within dorsal anterior insula, or dAI as used in (20). For each network, a two-sample t-test between superagers and typical older adults revealed that superagers had stronger intrinsic connectivity between the network seed and its targets (Fig. 2; Fig. 5B). Within the DMN, superagers showed stronger connectivity from the right PCC seed to right HF (Fig. 5B) as well as to a number of cortical targets, such as left superior frontal gyrus (SFG), right anterior middle temporal gyrus (aMTG), right ventrolateral prefrontal cortex (vlPFC), right pregenual anterior cingulate cortex (pgACC), right subgenual anterior cingulate cortex (sgACC) as well as bilateral angular gyrus (AG), dorsomedial prefrontal cortex (dmPFC), and rostromedial prefrontal cortex (rmPFC). Within the SN, superagers showed stronger intrinsic connectivity from the right dAI seed to the right supramarginal gyrus (SMG) and bilateral anterior midcingulate cortex (MCC) (Fig. 2). Furthermore, two-tailed planned contrast tests verified that superagers had significantly stronger connectivity when compared to typical older adults (*p* < 0.05; except left dmPFC, *p* = 0.051), whereas their intrinsic connectivity strength was statistically indistinguishable from young adults (*p* > 0.05; Fig. 3). Prompted by a reviewer’s comment, we conducted a post-hoc analysis examining connectivity between the DMN and SN (Fig. S2A). We found that superagers and young adults demonstrated similar connectivity between the right PCC (DMN node) and bilateral MCC (SN node) (p > 0.05), and that the inverse (negative) connectivity in young adults and superagers was not seen in typical older adults (p < 0.05) (Fig. S2B; see more details on this post-hoc analysis in SI). Notably, two-sample t-tests between superagers and typical older adults using control seeds in the motor and visual networks showed no statistical difference in intrinsic connectivity (motor: *t*(38) = 0.06, *p* = 0.95; visual: *t*(38) = -1.43, *p* = 0.16; Fig. 3).

**Figure 2.**
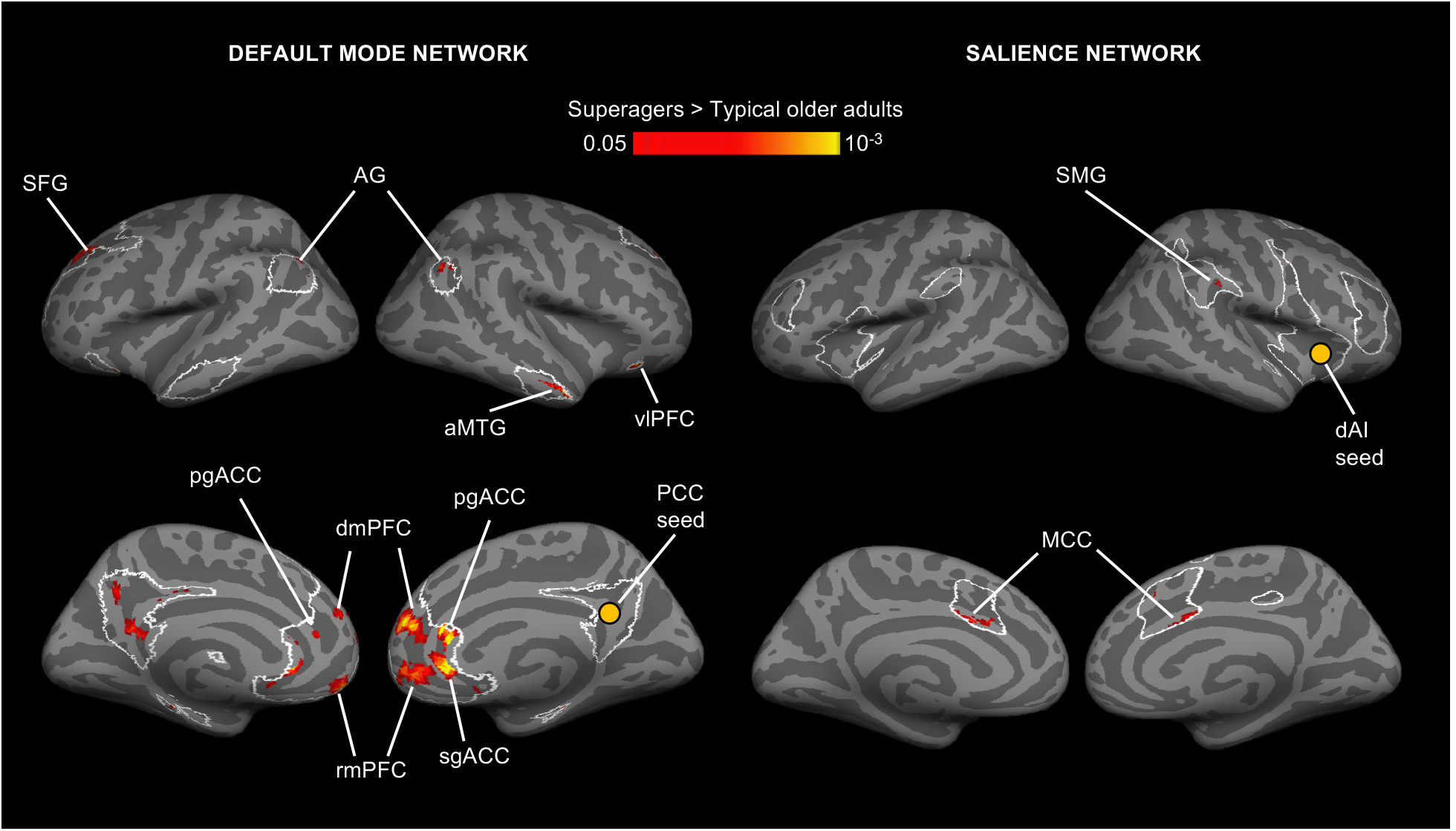
Regions of the DMN and SN (outlined in white) where superagers had stronger intrinsic functional connectivity than did typical older adults (red/yellow). For each network, a two-sample t-test between superagers and typical older adults was conducted. Maps were thresholded at *p* < 0.05 and masked by network masks shown in Fig. 1. We did not observe any region with higher connectivity in typical older adults than in superagers. We excluded clusters in the left precuneus/PCC region from further analyses because they showed high auto-correlation. AG, angular gyrus; aMTG, anterior middle temporal gyrus; dmPFC, dorsomedial prefrontal cortex; MCC, midcingulate cortex; pgACC, pregenual anterior cingulate cortex; rmPFC, rostromedial prefrontal cortex; SFG, superior frontal gyrus; sgACC, subgenual anterior cingulate cortex; SMG, supramarginal gyrus; vlPFC, ventrolateral prefrontal cortex.

**Figure 3.**
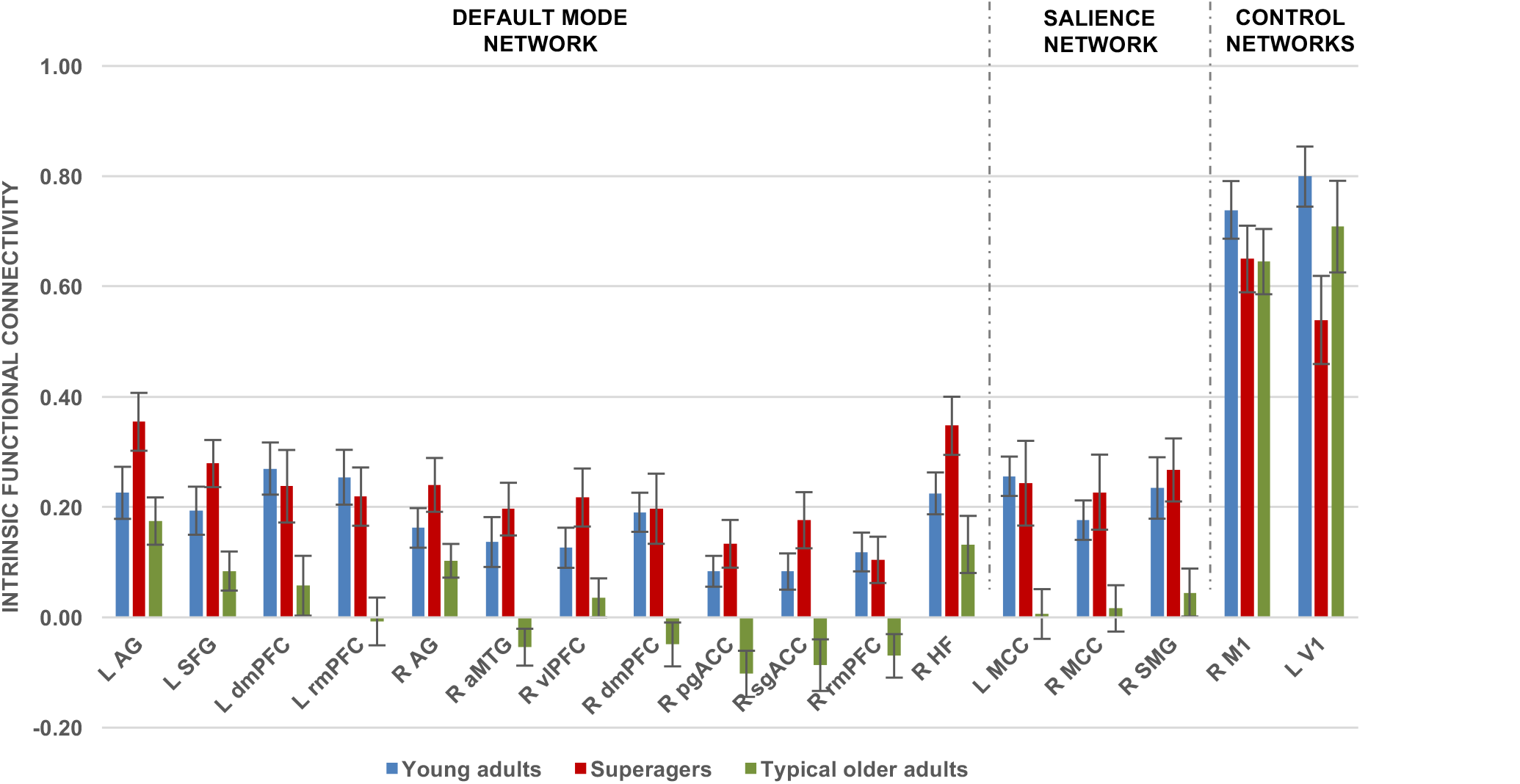
Preserved DMN and SN connectivity in superagers. Bar graphs show superagers had stronger intrinsic connectivity within DMN and SN than did typical older adults (*p* < 0.05) but similar connectivity to young adults. We calculated intrinsic connectivity strength between each network seed and its targets identified from peaks in the t-test maps in Fig. 2. These effects were specific to DMN and SN, as superagers did not differ from typical older adults in terms of visual or motor network connectivity strength. R, right hemisphere; L, left hemisphere. Error bars indicate 1 standard error of the mean. In several comparisons, the bars for superagers (orange) were higher than those for young adults (blue) and the error bars did not overlap. However, statistically, superagers were indistinguishable from young adults in all comparisons.

### Youthful Recall and Recognition Memory Performance in Superaging

We examined the generalizability of preserved memory in superagers using a visual-verbal paired-associates recognition memory task. Two one-way ANOVAs showed that the three groups differed on their item recognition memory [*F*(2,77) = 3.26, *p* < 0.05] and associative recognition memory [*F*(2,77) = 4.41, *p* < 0.05]. Planned contrasts indicated, as predicted, that superagers performed better than typical older adults in item recognition [*t*(77) = 2.04, *p* < 0.05, one-tailed] and marginally better in associative recognition [*t*(77) =1.44, *p* = 0.08, one-tailed] but did not differ significantly from young adults [item recognition: *t*(77) = -0.11, *p* = 0.92, two-tailed; associative recognition: *t*(77) = 1.10, *p* = 0.27, two-tailed]. To estimate the effect size of the difference between superagers and typical older adults, we computed Cohen’s d for each measure. Item recognition showed a large difference (*d* = 0.80, 95%CI = 0.49 -1.08) while associative recognition showed a moderate difference (*d* = 0.48, 95%CI = 0.08 -0.72). For comparison, CVLT free recall—used to define the groups—showed a large difference: *d* = 2.37, 95%CI = 1.96 - 3.26.

### Intrinsic Connectivity Predicted Recall and Recognition Memory Performance

We ran a series of bivariate correlation analyses between all three memory measures (CVLT verbal recall, visual-verbal item recognition, and visual-verbal associative recognition) and connectivity strengths of all 15 seed-target pairs (including both DMN and SN). As predicted, intrinsic connectivity strength of all seed-target pairs predicted recall performance in elderly adults at a False Discovery Rate (*q*) of 0.05 (Table 1; scatterplots for key nodes are depicted in Fig. 4A, B and Fig. 5C). Importantly, this brain-behavior relationship generalized to item recognition memory as well. Item recognition memory was significantly correlated with multiple indices of DMN connectivity strength (FDR-corrected at *q* = 0.05; Table 1). Additionally, a trend-level relationship was observed between item recognition memory and all three indices of SN connectivity strength, as well as between associative recognition memory and DMN network strength (Table 1). There was no association between memory performance and strength of intrinsic connectivity within the sensorimotor networks used as controls (Table 1). Post-hoc analysis of between-network connectivity revealed that connectivity (stronger inverse correlation) between right PCC and left MCC predicted better performance on CVLT (Fig. 4C), item recognition, and associative recognition; connectivity between right PCC and right MCC predicted only CVLT (Table S1).

**Table 1.** Association between memory and intrinsic connectivity within DMN and SN

| Target region | Recall |  | Item recognition |  | Associative recognition |  |
| --- | --- | --- | --- | --- | --- | --- |
|  | *r* | *p* | *r* | *p* | *r* | *p* |
| Default mode network |  |  |  |  |  |  |
| L AG | 0.44 | 0.00<sup>*</sup> | 0.17 | 0.15 | 0.23 | 0.08 |
| L SFG | 0.34 | 0.02<sup>*</sup> | -0.03 | 0.43 | 0.07 | 0.33 |
| L dmPFC | 0.34 | 0.02<sup>*</sup> | 0.22 | 0.09 | 0.37 | 0.01<sup>†</sup> |
| L rmPFC | 0.46 | 0.00<sup>*</sup> | 0.12 | 0.23 | 0.29 | 0.04 |
| R AG | 0.39 | 0.01<sup>*</sup> | 0.49 | 0.00<sup>*</sup> | 0.36 | 0.01<sup>†</sup> |
| R aMTG | 0.47 | 0.00<sup>*</sup> | 0.41 | 0.01<sup>*</sup> | 0.32 | 0.02<sup>†</sup> |
| R vlPFC | 0.40 | 0.01<sup>*</sup> | 0.43 | 0.00<sup>*</sup> | 0.23 | 0.08 |
| R dmPFC | 0.34 | 0.02<sup>*</sup> | 0.20 | 0.12 | 0.12 | 0.24 |
| R pgACC | 0.41 | 0.01<sup>*</sup> | 0.30 | 0.03<sup>†</sup> | 0.19 | 0.13 |
| R sgACC | 0.32 | 0.03<sup>*</sup> | 0.26 | 0.06<sup>†</sup> | 0.16 | 0.17 |
| R rmPFC | 0.38 | 0.01<sup>*</sup> | 0.27 | 0.05<sup>†</sup> | 0.12 | 0.23 |
| R HF | 0.34 | 0.02<sup>*</sup> | 0.27 | 0.05<sup>†</sup> | 0.32 | 0.02<sup>†</sup> |
| Salience network |  |  |  |  |  |  |
| L MCC | 0.43 | 0.00<sup>*</sup> | 0.28 | 0.04<sup>†</sup> | 0.09 | 0.30 |
| R MCC | 0.39 | 0.01<sup>*</sup> | 0.31 | 0.03<sup>†</sup> | 0.20 | 0.11 |
| R SMG | 0.38 | 0.01<sup>*</sup> | 0.29 | 0.04<sup>†</sup> | 0.22 | 0.09 |
| Control networks |  |  |  |  |  |  |
| R M1 | -0.08 | 0.31 | 0.09 | 0.30 | 0.01 | 0.48 |
| R V1 | 0.04 | 0.40 | -0.01 | 0.47 | 0.07 | 0.34 |

Note: *r* = Pearson's correlation coefficients, *p* = one-tailed significance. FDR-corrected significance: <sup>\*</sup>*q* = 0.05, <sup>†</sup>*q* = 0.10.

**Figure 4.**
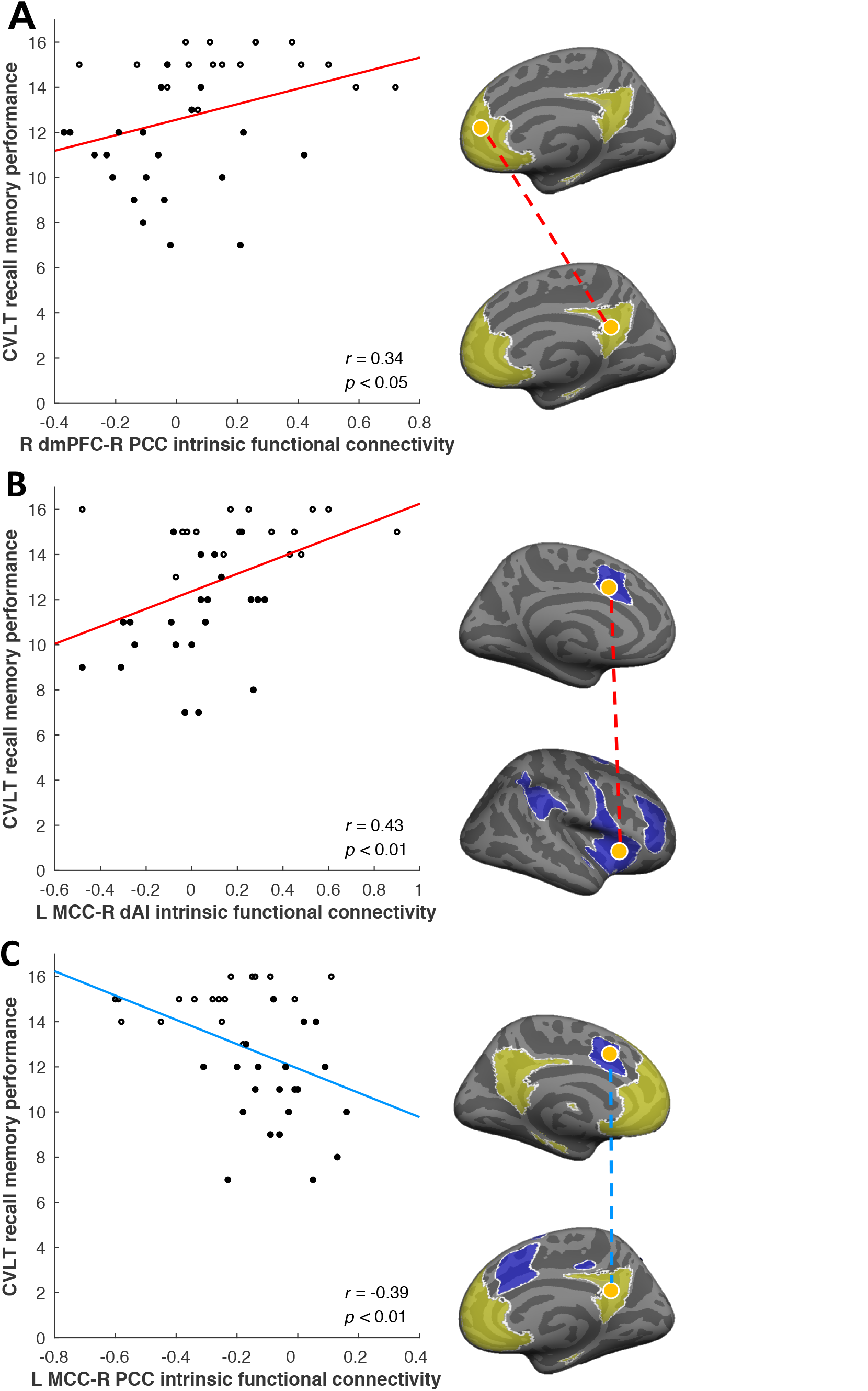
Preserved functional connectivity within and between default mode and salience networks supports preserved memory in elderly adults. Scatterplots illustrate the correlation between recall memory performance in the entire older adult group and intrinsic functional connectivity A) within the DMN (R dmPFC-R PCC), B) within the SN (L MCC-R dAI), and C) between the DMN and SN (L MCC-R PCC). Superagers are indicated by hollow points. The brain maps highlight the hypothesized networks of interest (DMN in yellow and SN in blue), seed and target nodes (in orange), and the direction of correlation representing functional connectivity (direct/positive in red and inverse/negative in light blue). Recall memory was scored out of a total of 16. Network seeds were R PCC (36) and R dAI (20). Displayed p-values are uncorrected.

**Figure 5.**
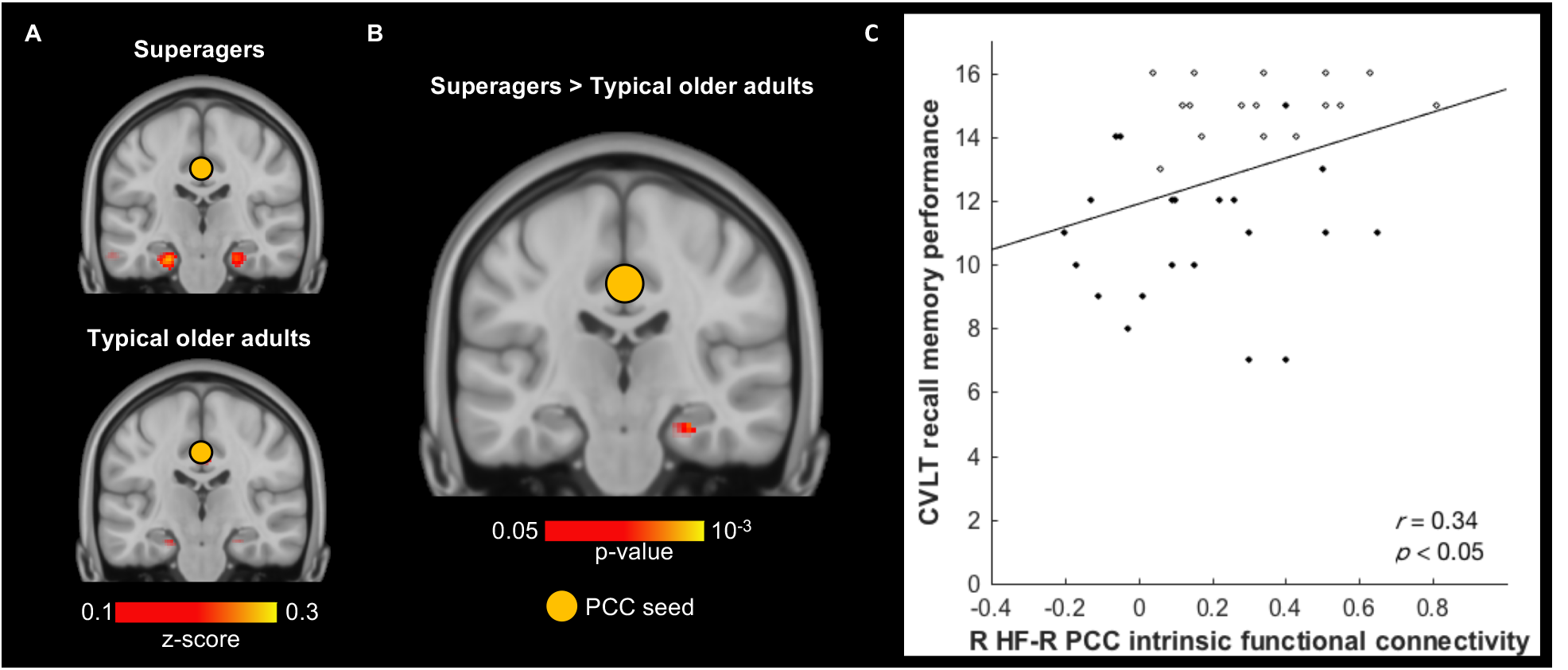
Preserved functional connectivity between hippocampus – a key node in the default mode network – and PCC correlates with preserved memory in elderly adults. A) Hippocampal connectivity to PCC seed (indicated by orange circle) in both elderly groups (*z* > 0.1). B) Two-sample t-test revealed that superagers had significantly stronger R HF-R PCC connectivity than typical older adults (*p* < 0.05). Hippocampal slices in both A and B were taken at y = -22 and displayed in neurological convention. C) R HF-R PCC connectivity positively predicted recall memory performance in the entire elderly group.

To test whether DMN and SN intrinsic connectivity each explained unique variance in memory performance, we ran one hierarchical linear regression analysis for each index of memory as the dependent measure, using intrinsic connectivity strength between canonical network nodes as predictor variables. We picked key nodes within the DMN and SN and used their connectivity strength as predictors in each regression model. For the DMN, the predictor was PCC connectivity to the right HF; for the SN, the predictor was dAI connectivity to the left MCC connectivity. We conducted two regression analyses, one using recall as the outcome measure and the other using item recognition as outcome measure. We did not conduct an analysis for associative recognition since it was not significantly predicted by connectivity between the dAI seed and any SN target. In both analyses, we observed that stronger intrinsic DMN connectivity and stronger SN connectivity *independently* predicted better memory performance, accounting for a total of 28% of the variance in the CVLT recall score (Table 2) and 15% of the variance in item recognition score (Table S2). Furthermore, between-network connectivity (L MCC-R PCC connectivity) only uniquely predicted item recognition score (Table S3), not CVLT recall score.

**Table 2.** Stronger functional connectivity in the DMN and in the SN independently predicted better recall memory in the elderly group.

|  | Recall memory performance |  |  |  |
| --- | --- | --- | --- | --- |
|  | *B* | *t* | *R*<sup>2</sup> change | Total *R*<sup>2</sup> |
| R HF-R PCC connectivity<sup>a</sup> | 0.31 | 2.19 | 0.10 | 0.28 |
| L MCC-R dAI connectivity<sup>b</sup> | 0.41 | 2.85 | 0.17 |  |

Note: This table displays standardized regression coefficient (*B*), *t* statistic (*t*), incremental variance (*R*<sup>2</sup> change), as well as total variance (Total *R*<sup>2</sup>) in recall memory performance predicted by entering both independent variables into a single multiple linear regression model. Incremental *R*<sup>2</sup> change indicates additional variance explained by respective independent variable when entered last in the model. <sup>a</sup>*p* = 0.035, <sup>b</sup>*p* = 0.007.

### Relationship between Neuroanatomy, Intrinsic Connectivity and Memory Performance

We additionally confirmed that for each network, intrinsic connectivity independently predicted recall memory (CVLT) over and above the anatomical preservation of network regions, as reported in Table 3. To index anatomical preservation, we initially planned to include cortical thickness/volume estimates for the regions used to compute estimates of intrinsic connectivity (i.e. R HF and R PCC for the DMN, L MCC and R dAI for the SN). R PCC thickness was not preserved in superagers (16) and R dAI thickness was not independently associated with CVLT performance over and above R HF volume and L MCC thickness, and so we excluded R PCC and R dAI from the analyses. As a consequence, we ran two regression analyses to predict CVLT performance, one using R HF volume and R HF-R PCC connectivity as DMN predictors, and the other using L MCC thickness and L MCC-R dAI connectivity as SN predictors, respectively. In our analysis, DMN neuroanatomy and intrinsic connectivity accounted for 27% of the variance in CVLT performance (first data row of Table 3). SN neuroanatomy and functional connectivity accounted for 26% of the variance in CVLT performance (second data row of Table 3). Then we ran one additional regression analysis with anatomical and functional variables of both networks as predictors, we found that they accounted for a total of 44% of the variance in CVLT performance (Table S4). In all cases tested above, adding between-network connectivity index (L MCC-R PCC connectivity) did not significantly improve the regression models. Cortical thickness measures did not significantly predict item recognition memory or associative recognition memory so additional regression analyses were not necessary in that regard.

**Table 3.** Within DMN and SN, higher structural integrity and stronger intrinsic connectivity independently predicted better recall memory in elderly participants.

|  | Recall memory performance |  |  |  |
| --- | --- | --- | --- | --- |
|  | *B* | *t* | *R*<sup>2</sup> change | Total *R*<sup>2</sup> |
| Default mode network |  |  |  |  |
| Adjusted R hippocampal volume<sup>a</sup> | 0.40 | 2.73 | 0.16 | 0.27 |
| R HF-R PCC connectivity<sup>b</sup> | 0.30 | 2.08 | 0.09 |  |
| Salience network |  |  |  |  |
| Adjusted L MCC thickness<sup>c</sup> | 0.29 | 1.92 | 0.08 | 0.26 |
| L MCC-R dAI connectivity<sup>d</sup> | 0.35 | 2.34 | 0.12 |  |

Note: This table displays standardized regression coefficient (*B*), *t* statistic (*t*), incremental variance (*R*<sup>2</sup> change), as well as total variance (Total *R*<sup>2</sup>) in recall memory performance predicted by entering both independent variables into a single multiple linear regression model. Incremental *R*<sup>2</sup> change indicates additional variance explained by respective independent variable when entered last in the model. <sup>a</sup>*p* = 0.010, <sup>b</sup>*p* = 0.045, <sup>c</sup>*p* = 0.063, <sup>d</sup>*p* = 0.025.

## DISCUSSION

An increasing number of studies document that age-related cognitive decline is not inevitable, with some elderly adults maintaining such cognitive abilities as attention, verbal memory, visuospatial memory, visuospatial perception, mental math and reasoning (5-9, 11); this has been called ‘successful aging’ (60-62), ‘cognitive reserve’ (63), and ‘brain maintenance’ (64). In this study, we add to this growing body of work by showing that superagers exhibit preserved intrinsic connectivity in two core intrinsic brain networks - the DMN and SN – and this connectivity predicted both recall and recognition memory. These findings allowed us to more precisely characterize superagers’ remarkable memory abilities. Performance on tests of free verbal memory recall, such as the CVLT, is thought to involve largely recollection-based memory processes (65-68), as well as controlled search/retrieval processes. In contrast, the ability to correctly identify individual items during a recognition memory test is believed to rely primarily on familiarity-based memory processes (69-71). Our own and others’ prior research (12, 16) demonstrated that superagers have superior free verbal recall performance, suggesting that recollection-based memory processes are superior in them relative to typical older adults. Here we extend this to demonstrate that superagers have youthful item-level recognition memory test performance, therefore suggesting that their familiarity-based memory processes are also better than those of typical older adults. Furthermore, our findings suggest that the association between intrinsic connectivity and memory performance extends generally to both recollection (measured by the CVLT) and familiarity (item recognition memory), and extends across verbal and visual domains. This relationship was strongest within the DMN, although associations between SN connectivity and item recognition memory exhibited similar trends as well. Further studies with larger sample sizes are needed to fully characterize the relationship between intrinsic connectivity and associative recognition memory. Overall, these findings suggest that preserved network integrity in superagers provides a process- and domain-general memory advantage.

### Intrinsic Functional Integrity as a Neurobiological Substrate for Superaging

Prior to this study, histologic (15) and morphometric evidence (12, 13, 15, 16) provided a static snapshot of superagers’ neuroanatomy. The present findings demonstrate youthful DMN and SN connectivity in superagers which, in turn, was associated with better memory performance, consistent with studies showing that intrinsic connectivity between DMN nodes (e.g. PCC, HF, mPFC) (46, 47, 56, 57, 72) and between SN nodes (e.g. AI, pgACC) (46, 47, 56, 57, 72) predicts memory performance in young adults and in older adults (73). We also showed preliminary evidence that superagers had preserved inverse correlations between DMN and SN nodes, comparable to young adults. This is consistent with previous studies showing inverse correlations between intrinsic DMN and SN signals in young adults (74, 75). As previously reported (51), this inverse correlation diminishes in typical older adults, but is remarkably preserved in superagers. Importantly, we found that within all older adults, stronger inverse between-network connectivity was associated with better memory performance, consistent with previous findings that between-network connectivity predicts working memory capacity (76) and verbal skills (77). Taken together, these findings suggest that superior memory performance in aging is associated with stronger within-network coupling in the DMN and SN, as well as stronger inverse correlations between these two key networks.

The network seeds and targets that we used in the current study (chosen from prior published reports) partially overlap with the regions that we previously identified based on preserved neuroanatomy alone (16) (Fig. 6). Notably, there were several DMN regions whose cortical thicknesses did not differ between superagers and typical older adults, even though their connectivity profiles did. These include the PCC seed itself, vlPFC, pgACC, and sgACC. On the other hand, there were several SN regions whose connectivity profiles did not differ between superagers and typical older adults, even as their cortical thicknesses did. These include the MI, dlPFC, and IFG. These findings indicate that reduced cortical thickness or subcortical volume do not necessarily imply reduced intrinsic connectivity (or vice versa), so that both structural and functional variation may independently predict unique behavioral variation in memory performance.

**Figure 6.**
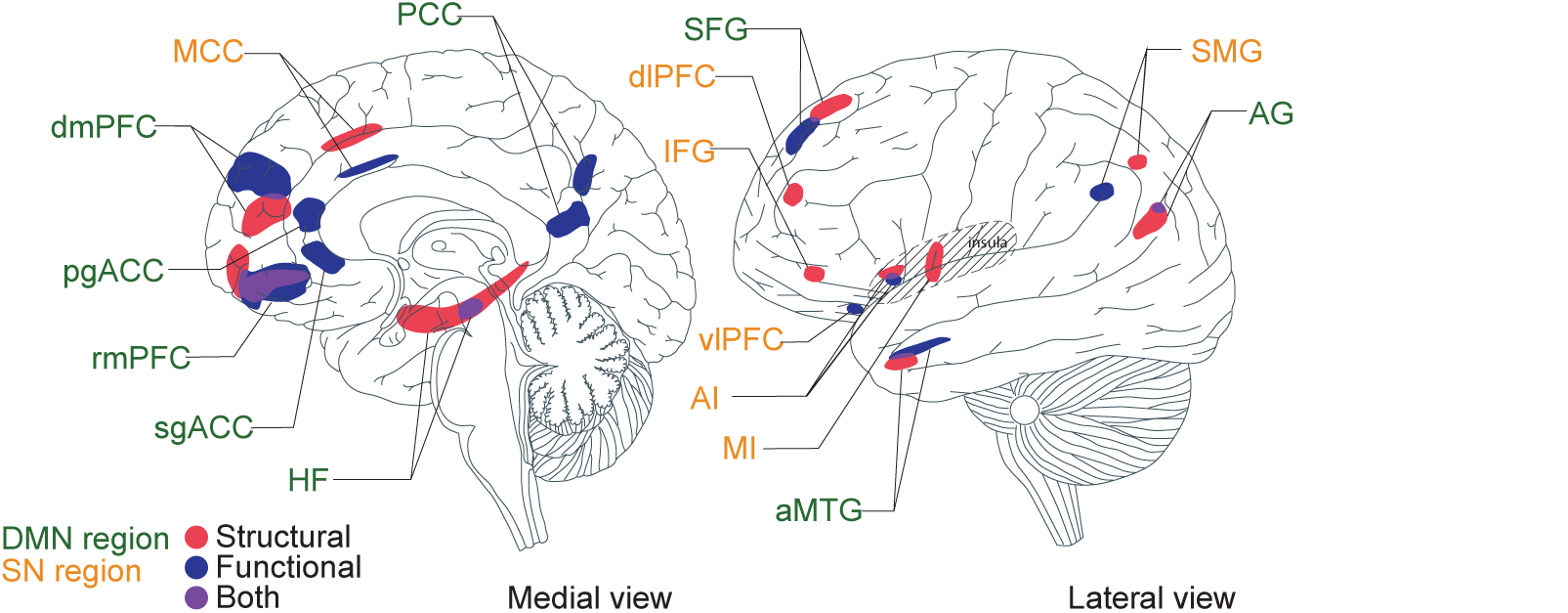
Structural and functional substrates of the superaging brain. Structural substrates (red) were defined based on preserved cortical thickness (16) and functional substrates (blue) were defined based on intrinsic functional connectivity. Overlapping regions are depicted in purple. Green font indicates DMN regions and orange font indicates SN regions. This is a summary schematic of all substrates combined into one hemisphere of the brain, with brain template based on (78).

### Limitations and Future Directions

The current study, with a sample size of 17 superagers, was limited in its power to detect small effect sizes. We are planning a future study to recruit a larger sample of superagers. Future studies might more comprehensively characterize functional connectivity for all observable resting state networks using methods such as the full correlation matrix analysis (79). Future research might also investigate the neural activation and dynamic connectivity while superagers are performing tasks.

It will also be necessary to determine exactly what functional role that DMN and SN play in superaging. A traditional interpretation of the role of DMN and SN in memory focuses on their preferential engagement in mnemonic and attentional or executive processes (20, 46). More recently, the DMN and SN have also been shown to be important for maintaining physiological balance (called “allostasis”, defined as anticipation and preparation to satisfy bodily needs before they arise; (80)) as well as interoception (awareness of, and sensitivity to, internal physiological sensations that result from allostasis; (81, 82). These findings suggest the hypothesis that superagers may better maintain allostasis in face of cognitive demand.

## MATERIALS AND METHODS

### Participants and Study Procedure

All participants (*n* = 91, 48 males) were right-handed native English speakers with normal or corrected-to-normal vision and none reported a history of substance neurological or psychiatric disorder. Ten participants were disqualified due to incomplete study procedure, resulting in a final sample size of 41 young adults (20 males, 24.5 ± 3.6 years) and 40 older adults (17 males, 66.9 ± 5.5 years). Participants completed the California Verbal Learning Test (CVLT) (83), the Trail Making Test (84), and associative memory task (72), structural and resting state scans. See Supporting Information for task details. As in our previous study (16), we defined superagers (*n* = 17, 5 males, 67.8 ± 6.0 years) as those who performed at or above the mean for young adults on the long delay free recall measure of the CVLT and no lower than one standard deviation below the mean for their age group on the TMT Part B. The remaining were classified as typical older adult (*n* = 23, 15 males, 66.2 ± 5.1 years). A list of all measures and tasks the participants completed can be found in Table S5. Demographic information, memory task data, group level connectivity maps, t-test maps, network masks and connectivity strength data can be retrieved on Open Science Framework (doi:10.17605/OSF.IO/G6F2K; ark:c7605/osf.io/g6f2k). Raw data and analysis scripts are also available upon request.

### MRI and fMRI

Structural data were acquired using a T1-weighted MPRAGE sequence (TR/TE/FA = 2530 ms/3.48 ms/7°; resolution = 1.0 mm isotropic). We reconstructed the cortical surface using the automated algorithm in Free-Surfer (http://surfer.nmr.mgh.harvard.edu) and extracted cortical thickness/subcortical volume from superaging brain regions as discussed in (16). Resting state data were acquired using an echo-planar sequence (TR/TE/FA = 5000 ms/30 ms/90°; resolution = 2.0 mm isotropic; 6.40 minutes, 76.8 volumes). To preprocess the resting state data, we removed first 4 volumes, corrected slice timing, corrected head motion, normalized to the MNI152 template, resampled to 2mm cubic voxels, removed frequencies higher than 0.08Hz, smoothed with a 6mm fwhm kernel and did nuisance regression (6 motion parameters, average global signal, average ventricular and white matter signals) (85-87).

### Functional Connectivity and Behavioral Correlation Analyses

We generated 4mm spherical regions of interest (ROIs) in the left posterior cingulate cortex (PCC; MNI 1, -55, 17) (36) to identify the DMN and in the right dorsal anterior insula (dAI; MNI 36, 21, 1) (20) to identify the SN. We computed Pearson’s *r* between the mean ROI time course and the rest of the brain, applied Fisher’s *r*-to-*z* transformation, and averaged the resulting maps across all subjects. For each network, we conducted a voxel-wise two-sample t-test between superagers and typical older adults. We identified peaks within masked t-test maps (Table S6) and extracted mean time courses within 4mm spherical ROIs. We calculated Pearson’s *r* between seed and all target time courses and applied Fisher’s r-to-z transformation. We calculated Pearson’s *r* between each network target’s connectivity strength and the 3 memory task scores (recall, item recognition and associative recognition), and used the Benjamini-Hochberg procedure (88) to correct for multiple comparisons. Control analysis was done using a motor network seed (M1; MNI -43, -16, 42) (89) and a visual network seed (V1; MNI -19, -95, 2) (90). Finally, we ran several hierarchical linear regression analyses to test how much variance in memory is explained uniquely by each network’s functional connectivity and neuroanatomy.

## ACKNOWLEDGEMENTS

This work was funded by Grant R01 AG030311 from the National Institute on Aging to L.F.B. and B.C.D.

